# mIDEA: An Interpretable Structure–Sequence Model for Methylation-Dependent Protein–DNA Binding Sensitivity

**DOI:** 10.1101/2025.11.14.688575

**Authors:** Yafan Zhang, Rina Li, Junhao Zhong, Xingcheng Lin

## Abstract

DNA methylation dynamically reshapes protein–DNA interaction landscapes, yet the mechanistic principles governing methylation-dependent recognition remain incompletely understood. Currently, there is no predictive framework that integrates methylation-aware binding specificity with 3D structure-based residue interactions for quantitatively assessing the impact of DNA methylation on protein–DNA interactions. To bridge this gap, we introduce mIDEA (Methylation-informed Interpretable protein–DNA Energy Associative model), a structure-aware, residue-level biophysical framework that predicts and interprets protein–methylated-DNA interactions by optimizing an amino acid–nucleotide energy matrix explicitly accounting for the modulatory effect of 5-methylcytosine. We validate mIDEA across diverse human transcription factors with distinct CpG methylation preferences. Using only structural templates and prior biochemical knowledge, mIDEA accurately recapitulates both “methyl-plus” and “methyl-minus” effects of DNA methylation on protein–DNA binding. Incorporating quantitative methylation-sensitive binding data further enables mIDEA to capture position-specific methylation effects. When integrated with whole-genome methylation profiles, the model enhances in vivo binding-site prediction by reducing false positives while preserving sensitivity compared to models trained on unmethylated sequences. Overall, mIDEA provides a principled framework for elucidating the molecular mechanisms underlying methylation-modulated protein–DNA recognition. The mIDEA source code is available at: https://github.com/LinResearchGroup-NCSU/IDEA_DNA_Methylation.

## 1 Introduction

5-methylcytosine (5mC) is a cytosine modified with a methyl group at the C5 position and occurs predominantly at CpG dinucleotide sites. Although this modification is chemically simple, it is one of the most prevalent DNA epigenetic marks across all kingdoms of life[1,2], including bacteria, plants, and mammals. 5mC plays fundamental roles in gene regulation[3,4,5,6], development[7,8,9], and cellular identity[10,11]. One major mechanism by which 5mC exerts its function is through altering the structural properties of DNA, which in turn reshapes protein–DNA binding landscapes[12,13,14]. For example, CpG methylation can repress transcription by preventing the binding of certain transcription factors, whereas some proteins specifically recognize 5mC and may trigger local demethylation[6,15,9], thereby contributing to cellular reprogramming and differentiation. Understanding how cytosine methylation modulates protein-DNA recognition[16,17,18,19,20] is therefore essential for elucidating the regulatory logic underlying gene expression and epigenetic control.

Traditionally, classic low-throughput methods[12] such as electrophoretic mobility shift assays (EMSA), isothermal titration calorimetry (ITC), and DNase I footprinting have been used to accurately and quantitatively measure protein–DNA interactions involving methylated DNA, offering precise control over oligonucleotide design. More recently, several high-throughput technologies, including EpiSELEX-seq[21], methyl-HT-SELEX[22], methyl-Spec-seq[23], and methyl-PBM[24], have enabled the systematic profiling of transcription factor (TF) methyl-sensitivity at scales ranging from 10^4^ to 10^6^ methylated oligonucleotides in a single experiment. In particular, methyl-HT-SELEX has been applied to investigate the effects of 5mC on the binding of more than 500 human TFs, revealing distinct protein-specific preferences for CpG methylation (mCpG). For example, some developmentally important TFs exhibit a strong preference for binding to methylated CpG sites (“methyl-plus”), whereas others, such as MAX, preferentially bind unmethylated sites (“methyl-minus”). Interestingly, a small subset of TFs shows position-dependent methylation sensitivity within their binding motifs. Although these high-throughput approaches have greatly expanded our understanding of TF methylation-dependent binding specificity, the mechanistic basis underlying these differential preferences remains poorly understood.

With the advancement of experimental technologies, a variety of computational methods have been developed to model and quantify the effects of cytosine methylation on protein–DNA binding affinity and specificity. For example, Kribelbauer *et al*. developed a feature-based generalized linear model trained on EpiSELEX-seq data to estimate position-specific methylation effects. A more direct and widely used strategy is to extend the DNA alphabet to include 5-methylcytosine (5mC) as an additional “fifth” nucleotide, enabling the learning of methylation-aware binding models. For instance, ProBound[25] is a machine learning–based model that captures methylation effects and outputs absolute *K*_*d*_ values. Similarly, Viner *et al*.[26] expanded position weight matrices (PWMs) to incorporate methylated cytosines, revealing methylation-sensitive motifs. Other models, such as MeDeMo[27] and SEMplMe[28], use an extended six-letter alphabet with separate symbols for methylated cytosines and the corresponding guanines on the opposite strand, thereby capturing methylation-dependent specificity and intra-motif dependencies. Notably, unlike these sequence-based models, the chemistry-driven DeepRec framework represents DNA using physicochemical signatures. Trained on methylated HT-SELEX data, DeepRec[29] can predict the impact of mCpG on relative protein–DNA binding affinity and provide mechanistic insights by linking its learned energy logo to available structural data. Although these approaches have proven powerful for predicting methylation effects and discovering methylation-sensitive motifs, they typically require large amounts of high-throughput sequencing data for training. However, such data can be costly to generate and are often unavailable for many protein systems, limiting their broader applicability.

On the other hand, the growing availability of structural data[30], together with recent advances in AI-based structure prediction[31,32,33,34], has greatly expanded the repertoire of available 3D structures for protein–DNA complexes. Such an expansion enables the development of new predictive models[35,36] that incorporate protein–DNA complex structures during training, thereby reducing the dependence on high-throughput binding sequence data. Building on this concept, we previously developed a sequence-specific protein–DNA binding affinity model, IDEA[36], which leverages the principle of minimal frustration[37,38] and experimentally determined protein–DNA complex structures to optimize a physicochemical energy model for predicting sequence-specific protein–DNA binding affinities. The model demonstrated superior predictive accuracy while requiring substantially fewer training sequences.

In this manuscript, we extend the IDEA model to study DNA methylation by introducing an additional alphabet symbol for 5-methylcytosine (5mC), thereby developing the Methylation-informed Interpretable protein–DNA Energy Associative (mIDEA) model (Figure 1). The mIDEA framework integrates binding specificity data with experimental or AI-predicted structural information and learns an interpretable residue-level energy matrix that quantifies how cytosine methylation modulates protein–DNA binding affinities. In addition, mIDEA provides mechanistic insights into methylation-dependent recognition across proteins with distinct mCpG binding preferences. By further integrating whole-genome bisulfite sequencing (WGBS) data, we developed a genomic binding-site prediction pipeline that explicitly incorporates methylation profiles. Altogether, mIDEA establishes an integrative structure–sequence framework for predicting the modulatory effects of DNA methylation and elucidating its mechanistic role in shaping gene expression.

**Fig. 1.**
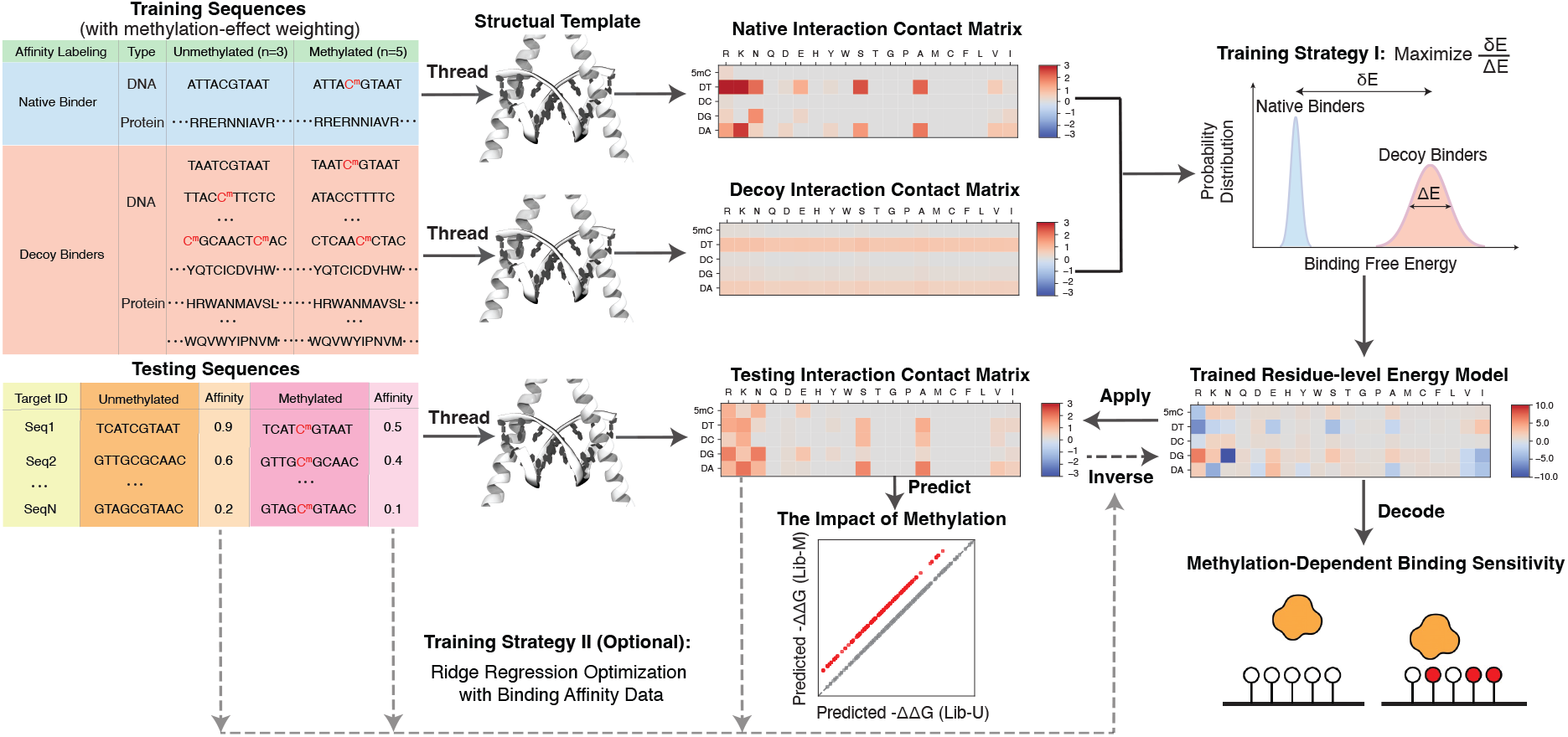
Overview of the mIDEA framework. **Input**: Training sequences include both native binders (derived from crystallized structures or identified by high-throughput experiments) and artificially generated decoy binders, each with methylated and unmethylated versions. Prior methylation-effect knowledge is incorporated by assigning copy-number weights to the methylated and unmethylated training sequences (e.g., 5:3 for the weak methylplus protein C/EBP*β*). Each training or testing sequence is threaded onto a structural template to compute its residue–nucleotide interaction contact matrix *ϕ*(*a, n*). **Process:** Two complementary training strategies are employed to optimize the residue-level energy model *γ*(*a, n*). (*i*) **Strategy I:** Maximizes the energy separation between native and decoy binders (*δE/ΔE*). (*ii*) **Strategy II:** When quantitative affinity data are available, ridge regression is applied to directly optimize *γ*(*a, n*) by minimizing the mean squared error between experimental and predicted binding energies. **Output:** The optimized residue-level energy model decodes the mechanistic basis of protein sensitivity to DNA methylation. Applying the trained model to the testing interaction contact matrices yields quantitative predictions of the impact of DNA methylation on protein–DNA binding.

## 2 Materials and Methods

### 2.1 mIDEA Framework

mIDEA (Figure 1) extends the original IDEA framework [36], a structure–sequence model for predicting sequence-specific protein–DNA binding affinities, by explicitly accounting for cytosine methylation. To achieve this, mIDEA introduces a “fifth” nucleotide type—5-methylcytosine (5mC)—into the energy formulation, thereby expanding the residue-level interaction matrix from 20 *×* 4 to 20 *×* 5 to represent interactions between amino acids and nucleotides. This design enables mIDEA to integrate structural information with binding-sequence data from both experimentally determined and AI-predicted protein–DNA complexes, as well as from high-throughput methylation-specific assays. The framework predicts the effects of cytosine methylation on protein–DNA binding affinity and provides residue-level mechanistic interpretations of methylation-dependent binding preferences. All datasets used for model training and evaluation are publicly available and summarized in Section S1.1.

#### 2.1.1 Protein-DNA Energy Model

mIDEA employs a coarse-grained, residue-level representation of protein-DNA interactions. The binding interface is defined by all pairs of protein C*α* atoms and DNA phosphate (P) atoms within an 8 Å distance cutoff. For a given complex, the residue-nucleotide interaction landscape is represented by an interaction contact matrix *ϕ*(*a, n*), formulated as a function of amino acid type *a* and nucleotide type *n* ∈ A, T, C, G, 5mC.

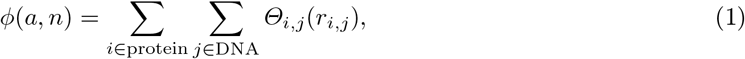

where the smooth contact function *Θ*_*i,j*_(*r*_*i,j*_) is given by:

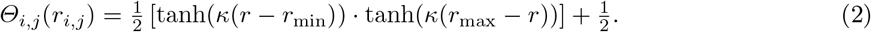

Here, *r*_min_ = −8Å and *r*_max_ = 8Å define the distance bounds, with *κ* = 0.7 controlling the transition steepness. This function asymptotically approaches 1 for close contacts (*r*_*ij*_ *<* 4 Å), decreases smoothly for 4 ≤ *r*_*ij*_ ≤ 8 Å, and converges to 0 for *r*_*ij*_ ≫ 8 Å, thereby defining a continuous and differentiable contact weight rather than a hard distance cutoff.

The total binding energy of a protein-DNA complex is a linear function of its contact matrix:

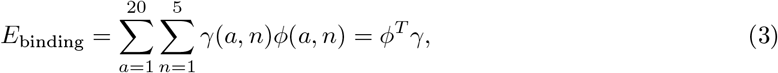

where *γ*(*a, n*) is the optimized 20 × 5 energy parameter matrix.

To predict binding affinity for a new protein-DNA sequence pair, the target sequence is threaded onto an appropriate structural template and its interaction contact matrix *ϕ*_target_ is computed. The predicted binding energy is then:

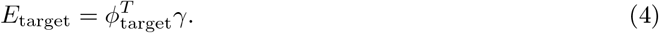

As *γ* is determined up to an arbitrary scaling factor, *E*_target_ is presented in reduced units, enabling comparison of relative binding affinities across different sequences.

Treating 5mC as a distinct nucleotide type enables mIDEA to explicitly capture the impact of cytosine methylation. The difference *γ*(*a*, 5mC) ™ *γ*(*a*, C) directly quantifies the residue-level energetic perturbation induced by methylation, providing an interpretable measure of methylation sensitivity. To quantify the impact of CpG methylation on binding affinity, we define:

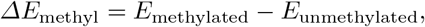

which represents the methylation-induced energetic shift for the same DNA sequence.

The thermodynamic relation [39] connects these energy differences to relative binding affinities:

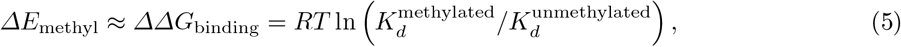

where *ΔE*_methyl_ can be converted to relative dissociation constants *ΔK*_*d*_, providing direct quantification of methylation’s effect on binding affinity.

In this study, we developed two complementary strategies for optimizing the interaction energy matrix *γ*(*a, n*), which together form the core of mIDEA’s predictive power. The first strategy integrates structural templates, synthetic sequence decoys, and prior knowledge of how cytosine methylation influences DNA interactions with target proteins to construct a sparse but comprehensive training dataset. This enables mIDEA to capture methylation effects even in the absence of detailed quantitative data. In contrast, the second strategy refines *γ*(*a, n*) in a fully data-driven manner by fitting the model directly to quantitative protein–methylated DNA binding measurements. By mapping experimental signals onto the geometric features of the binding interface, this approach provides a more direct representation of methylation-dependent interaction patterns. Together, these two strategies allow mIDEA to flexibly integrate both prior knowledge and experimental data, balancing interpretability with predictive accuracy.

#### 2.1.2 Strategy I: Optimizing Methyl-Binding Specificity via Prior Knowledge and Sequence Decoys

This strategy optimizes *γ*(*a, n*) using structural templates and prior knowledge about a protein’s general methylation response, without requiring quantitative binding data. The process begins by identifying a high-quality structural template (either experimental or AI-predicted) for a protein bound to a top-binding DNA sequence containing a CpG site. Both non-methylated and methylated versions of this top-binding sequence are then threaded to replace the existing DNA sequences within the template to form the training dataset.

Additionally, prior knowledge of the methylation effect is incorporated by assigning different weights for the training sequences. For instance, a weak methyl-plus effect is encoded by assigning 5: 3 weight favoring methylated over non-methylated interactions, while a strong effect uses a 10: 1 ratio. The prior knowledge was obtained from experimental observations of methylation-dependent binding effects, ranging from low-throughput methods[12] (EMSA, ITC, DNase I footprinting) to high-throughput assays such as EpiSELEX-seq[21], methyl-HT-SELEX[22], methyl-Spec-seq[23] and methyl-PBM[24].

The model is trained by maximizing the energetic separation between the training strong binders (the threaded top-binding sequences) and artificially generated decoy binders. Specifically, the decoys were generated as scrambled variants of the strong binders through base randomization that preserved length and nucleotide composition, producing 1,000 sequences by randomizing the DNA and 10,000 by randomizing the protein sequences. Similar strategies have demonstrated success in capturing sequencespecific effects in protein folding[37,38,40], protein-protein[41,42], and protein-nucleic acid interaction studies[36,43], especially when experimental binding data are scarce. The optimization objective aims to maximize the energy gap between strong and decoy binders, normalized by the standard deviation of the decoy binders:

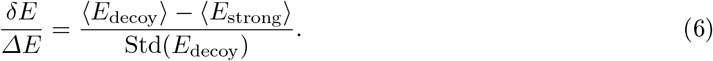

Building on the interaction contact matrix *ϕ*(*a, n*) defined in Eq. 1, we next define the feature differential vector and the decoy covariance matrix as:

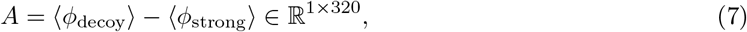

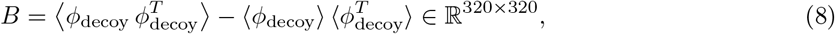

the objective function can be reformulated as 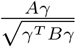, with the analytical solution being *γ* ∝ *B*^*−*1^*A*.

To ensure numerical stability and physical interpretability, we regularize the solution by:

1. Diagonalizing *B* = *PΛP* ^*−*1^;
2. Retaining the top 60 eigenmodes;
3. Replacing smaller eigenvalues with the 60th largest value;
4. Reconstructing *B*^*−*1^ from the filtered eigenmodes.

The resulting *γ* matrix can be reshaped to enable interpretations of residue-level methylation preferences.

#### 2.1.3 Strategy II: Integration of Detailed Specificity from Binding Data

When highthroughput, methylation-sensitive binding data (e.g., from EpiSELEX-seq) is available, *γ* can be optimized directly from quantitative measurements. Relative binding affinities are derived from foldenrichment profiles and converted into a 1-D array of experimental binding energies, *E*_exp_ ∈ ℝ^*N*^, where *E*_exp_ = *RT* ln *K*_*d*_ for each of the binding sequences.

Given the corresponding interaction features derived from the interaction contact matrix *ϕ* (*a, n*) defined in Eq. 1, we construct a feature matrix *Φ* ∈ ℝ^*N ×*320^ by stacking the *ϕ* vectors for all *N* DNA targets. We then solve for the interaction parameter vector *γ* using ridge regression:

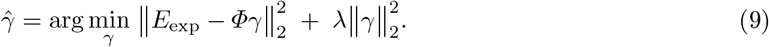

The optimal regularization strength *λ*^*^ is determined via five-fold cross-validation to minimize the mean squared error (MSE) on held-out validation sets, ensuring a balanced trade-off between bias and variance.

To ensure robustness, the optimization is repeated over 50 independent random train–test partitions, and the final model is refit using the averaged *λ*^*^ on the entire dataset. The MSE is defined as:

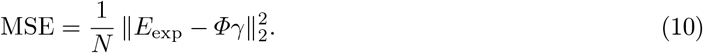

### 2.2 Methylation-Informed Genomic Protein–DNA Binding Sites Prediction

In this study, we took an initial step toward applying the mIDEA framework to predict and analyze how CpG methylation modulates the genomic protein–DNA binding landscape. The methodological details are described below.

#### 2.2.1 Quantification of CpG Methylation

To characterize the local methylation landscape, we quantified CpG density and methylation status across the genomic sequence using non-overlapping 1 kb bins. CpG dinucleotides were first identified and counted based on the GRCh38 reference genome. Wholegenome bisulfite sequencing (WGBS) data aligned to the same reference were then used to classify each CpG site as *unmethylated* (≤30%), *intermediate* (30–70%), or *methylated* (≥70%) according to the proportion of reads showing methylation at that locus. For each bin, the numbers of CpG sites in the three categories were aggregated and normalized by the total CpG count, yielding the relative fractions of unmethylated, intermediate, and methylated CpGs across the region.

#### 2.2.2 Computation of Methylation-Informed Energy Scores

We next incorporated local methylation information into the predicted binding energy landscape. First, we adopted a sliding-window approach in which the genomic sequence was segmented into overlapping fragments of the same length as the training sequence (e.g., 11 bp for MAX) from the 5^*′*^ to 3^*′*^ end in 1-nucleotide increment. The binding free energy for each fragment was then computed following the procedure described in Section 2.1.1. Two baseline free energy profiles were obtained: *E*_un_ represents the predicted energy values when all CpG sites within the genomic sequence are treated as unmethylated, whereas *E*_me_ represents the corresponding energies when all CpG sites are assumed to be fully methylated. Each energy profile was subsequently normalized to a *Z*-score using the following equation, yielding *Z*_un_ and *Z*_me_:

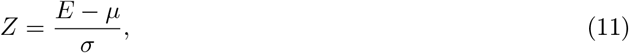

where *E* denotes the raw energy value, and *μ* and *σ* represent the mean and standard deviation of all energy values across both unmethylated and methylated conditions, respectively.

To account for the observed methylation heterogeneity, we derived a methylation-informed energy score (*Z*_informed_) by combining the unmethylated and methylated energy states according to their local CpG methylation fractions. The unmethylated (*r*_unmeth_), intermediate (*r*_interm_), and methylated (*r*_meth_) fractions were obtained from Section 2.2.1. The final energy score was computed as:

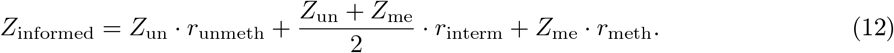

This formulation provides a continuous representation of methylation influence on the local binding energy landscape by proportionally weighting the energetic contributions of unmethylated and methylated states according to their observed CpG methylation levels.

## 3 Results

### 3.1 mIDEA Decodes Methyl-Plus Binding Sensitivity

As a proof of mIDEA’s utility, we first focused on the bZIP transcription factor C/EBP*β*, which has previously been reported to be a weak methyl-plus protein by both EpiSELEX-Seq[21] and methyl-HT-SELEX[22] (Figure S1). We aimed to evaluate whether our model could accurately predict such weak 5mC-dependent binding effects. For training, we used the co-crystal structure of C/EBP*β* bound to DNA (PDB ID: 8K8D; Figure 2A), which contains a high-affinity palindromic binding site (5’-CATTACGTAATG-3’). The central 10-mer sequence (5’-ATTACGTAAT-3’) corresponds to a top motif identified by EpiSELEX-seq. To model the weak methyl-plus specificity, we trained mIDEA using two DNA templates, following the procedure described in Section 2.1.2. Specifically, five copies of a sequence containing a methylated cytosine (5’-CATTA(5mC)GTAATG-3’) and three copies of its unmethylated counterpart (5’-CATTACGTAATG-3’) were provided. This 5:3 ratio of methylated to unmethylated templates corresponds to an approximate 1.7-fold enrichment, which is consistent with the 1.5–2× binding preference measured by EpiSELEX-seq. mIDEA successfully captured the methyl-plus effect of C/EBP*β*, showing a consistent upward energy shift for all CG-containing sequences (Figure 2B).

**Fig. 2.**
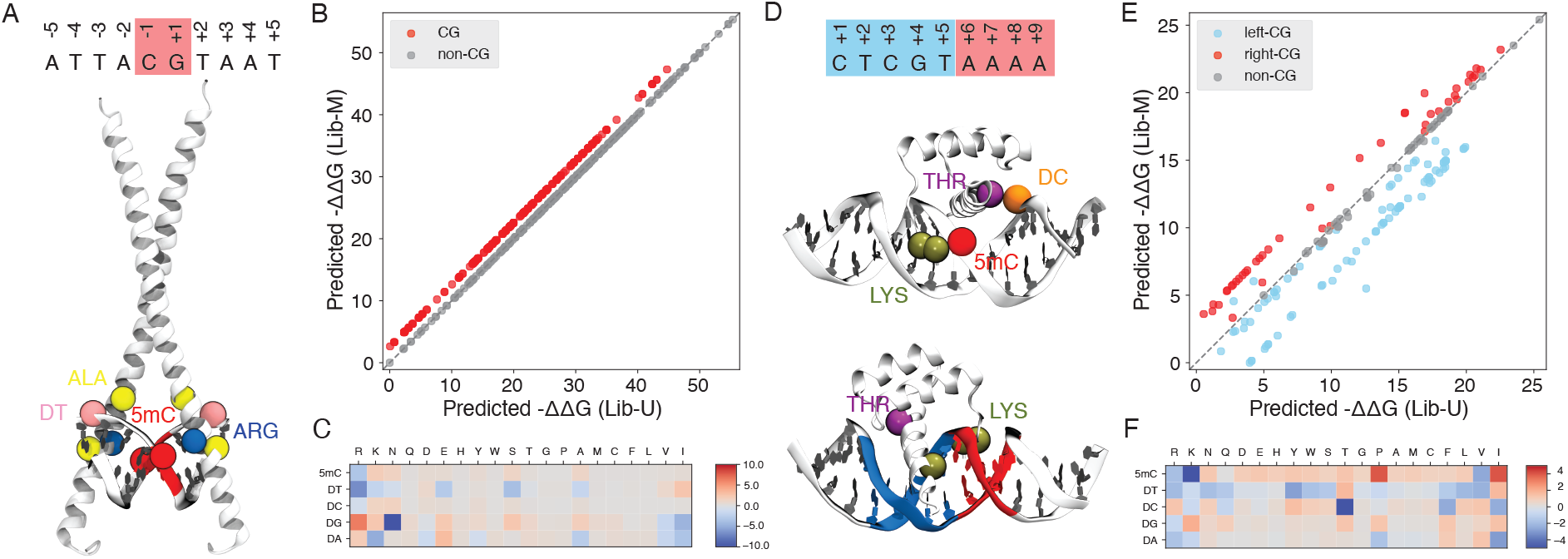
Modeling weak and strong methyl-plus binding effects using mIDEA. (**A**) Crystal structure of the C/EBP*β*–DNA complex (PDB ID: 8K8D), highlighting the central CG dinucleotide and key residues involved in methyl group recognition. (**B**) Binding free energy predictions (*Predicted ™ΔΔG/RT*) for C/EBP*β* on methylated (Lib-M) versus unmethylated (Lib-U) ligands. Each point represents a 10-mer sequence, with its position relative to the diagonal indicating the effect of methylation (points above the diagonal denote enhanced affinity). The overall upward shift of CG-containing ligands reveals a weak methyl-plus effect for C/EBP*β*. (**C**) Trained residue-level energy model for C/EBP*β*, where bluer shades indicate stronger interactions. The model captures a strong preference of ARG for both 5mC and DT, with a favorable ARG–5mC interaction underlying the weak methyl-dependent affinity increase. (**D**) Crystal structure of the HOXB13–methylated DNA complex (PDB ID: 5EF6), highlighting key residues and the right (red) and left (blue) mCpG regions used for structural interpretation. (**E**) Scatter plot of predicted binding free energies of HOXB13 on methylated (Lib-M) versus unmethylated (Lib-U) 9-mers. Red points (right-CG ligands) cluster above the diagonal, indicating strong methylation-enhanced binding, whereas blue points (left-CG ligands) show a markedly weaker effect, revealing a distinct position-dependent methyl-plus specificity. (**F**) Trained residue-level energy model for HOXB13, revealing strong LYS–5mC and THR–DC interactions consistent with the structural contacts.

The corresponding trained energy model (Figure 2C) highlights a strong interaction between the ARG residue and the cytosine base—both in its unmethylated (DC) and methylated (5mC) forms—with the 5mC interaction being energetically more favorable. This short-range ARG–5mC interaction provides a mechanistic explanation for the methylation-enhanced binding of C/EBP*β* to DNA. This finding is consistent with results from DeepRec [29], which was trained on EpiSELEX-seq data and further supported by their independent analysis of a C/EBP*β*–DNA co-crystal structure (PDB ID: 6MG2). In that study, DeepRec reported a favorable van der Waals interaction between an ALA residue and the methyl group of thymine—a feature that our model likewise captures. Notably, mIDEA does not learn such methyl-dependent contacts in isolation; it simultaneously acquires residue–nucleotide interaction patterns for both methylated and canonical bases, enabling a unified representation of methylation-specific and sequence-specific binding energetics. In particular, it identified a strong ARG–DT interaction, indicating the methyl group carried by both thymine and 5mC leads to their favorable interactions with ARG through similar physicochemical interactions.

To further assess whether our model can capture strong methyl-plus effects, we selected another example: the homeodomain protein HOXB13, which was previously reported to exhibit strong methylplus binding behavior in methyl-HT-SELEX[22]. We analyzed the Round 1 and Round 4 data from both the methyl-HT-SELEX and normal-SELEX experiments of the original study using the R package SELEX [44]. We analyzed the Round 1 and Round 4 data from both the methyl-HT-SELEX and normal-SELEX experiments using the R package SELEX [44]. Our 9-mer–based statistical analysis successfully reproduced the ligand-enrichment patterns reported by Yin et al. [22] (Figure S2A), showing that CG-containing sequences were 1–10 times more abundant in the methylated library (Lib-M) than in the unmethylated library (Lib-U). We further quantified the binding affinity (Figure S2B) of each 9-mer ligand based on its enrichment from the first to the fourth round, which revealed several key insights. First, methylation of the CG dinucleotide positioned on the right side of the motif generally enhanced the binding affinity for HOXB13. In contrast, methylation of the left-side CG exhibited variable effects. Second, although non-CG ligands were expected to cluster along the diagonal and reflect no major change in binding affinity, many of them exhibited an downward shift, suggesting that their effective binding was reduced in the methylated library (Lib-M), possibly due to increased competition from CG-containing ligands for HOXB13 binding. The SELEX data also revealed that HOXB13 primarily recognizes the motif 5’-CTCGTAAAA-3’. Methylation of the cytosine in this motif resulted in both higher ligand counts and stronger binding affinity. Collectively, our analysis confirms that HOXB13 exhibits a binding preference for the methylated version of its primary motif and for the majority of right-side CG-containing sequences. We trained the model on a crystallized HOXB13–DNA co-complex with a methylated core motif (PDB ID: 5EF6; Figure 2D), using a 10:1 ratio of methylated to unmethylated templates to reflect the tenfold increase in CG-ligand counts from Lib-U to Lib-M observed in the processed methyl-HT-SELEX data (Figure S2A). Consistent with the experimental observation (Figure S2B), the model did not show a uniform upward shift for all CG-containing sequences (Figure 2E), but rather a clear positional effect: HOXB13 has a distinct preference to bind methylated right-CG ligands over their unmethylated counterparts and the methylated left-CG ligands. This result supports that the model is not merely memorizing the training sequence templates—despite being trained on a methylated left-CG sequence—but instead captures meaningful right-CG methylation-enhanced effects beyond the input training data. To further investigate the mechanistic basis of this behavior, we examined the 3D structure of the HOXB13–methylated DNA complex (Figure 2D) and the trained energy model (Figure 2F). The model revealed that LYS interacts more strongly with 5mC than with unmethylated cytosine, whereas THR shows the opposite trend, exhibiting a stronger interaction with canonical cytosine than with 5mC. These findings are consistent with the close LYS–5mC and THR–DC contacts observed in the structural data (Figure 2D). Furthermore, the enhanced contribution of the right-flanking mCpG region can be attributed to the geometric arrangement, where LYS residues are positioned closer to the right (red) region than to the left (blue). Similarly, the close proximity of THR to the left blue region explains why our model predicts reduced binding affinity when the cytosine at this position is methylated. Nevertheless, the experimentally observed methylation effects for left-CG sequences show considerable variability (Figure S2B), and the underlying reason remains unclear. One possible explanation is that the left-CG sequences undergo greater conformational fluctuations, leading to larger fluctuations in binding affinities than the right-CG group. In addition, mIDEA revealed strongly favorable interaction energies between 5mC and the ARG and VAL residues, consistent with experimental findings that suggest residues with aliphatic chains contribute to favorable protein-DNA hydrophobic interactions [22].

### 3.2 mIDEA Decodes Methyl-Minus Binding Sensitivity

We next examined a group of transcription factors that display methyl-minus binding behavior, with MAX–a basic helix-loop-helix (bHLH) transcription factor–as a representative example. MAX specifically recognizes the E-box sequence 5’-CACGTG-3’ [45]. Based on methyl-HT-SELEX [22], CpG methylation of the E-box motif reduces its enrichment relative to the unmethylated sequence by 10–200 fold, accompanied by an approximately 25% decrease in binding affinity (Figure S3). Following the same training protocol in Section 2.1.2, we trained mIDEA using the MAX–DNA co-complex (PDB ID: 1HLO; Figure S4A), with a training set composed of a 10:1 ratio of unmethylated to methylated templates, thereby encoding strong methyl-minus specificity. The model successfully captured this effect, predicting reduced binding upon CpG methylation (Figure S4B).

The trained energy model further revealed that the interaction between ARG and normal cytosine is energetically more favorable than ARG–5mC interactions (Figure S4C), consistent with our previous training results[36], which were obtained using the same 1HLO structural template described above. In addition, GLN, SER, and ALA residues showed a slight preference toward unmethylated cytosine compared with 5mC. These unmethylated-cytosine–favored interactions likely underlie MAX’s methylminus specificity.

### 3.3 mIDEA Decodes Position-Specific Methylation Binding Sensitivity

Only a small subset (5%)[22] of transcription factors demonstrates position-dependent preferences for CpG methylation within their binding motifs. A representative example is the bZIP transcription factor ATF4, which binds the cyclic AMP response element (CRE motif, 5’-TGACGTCA-3’) as a homodimer. EpiSELEX-seq analysis (Figure S5) identifies multiple effects of CpG methylation on ATF4 binding. Methylation at flanking sites strengthens the interaction, whereas methylation at the central site weakens it. The flanking enhancement is the dominant factor, as shown by the fact that methylating both regions still increases affinity, although central CpG methylation partially offsets this gain and lowers it below the level produced by flanking methylation alone. Notably, the mechanistic basis for this position-specific effect of CpG methylation on ATF4 binding remains largely unknown[21,29].

Predictions and explanations of the position-dependent methylation effects of ATF4 binding are complicated by the absence of experimentally determined ATF4-DNA structures. To address this issue, we utilized a deep-learning-based structural prediction tool AlphaFold3 [31] to model a high-confidence ATF4-DNA complex (ipTM = 0.81, pTM = 0.82, Figure 3A), using the top binding motif containing the CRE motif from the EpiSELEX data [21]. As flanking methylation increases binding affinity by two-to ten-fold relative to the unmethylated state, we trained the mIDEA model using a 10:1 ratio of methylated to unmethylated left-flank CG sequences (5’-ACGATGTCAT-3’) to prioritize learning the dominant effect of flanking methylation. Under this structure-based training scheme, the model successfully captured both the enhancing effect of flank methylation and the suppressive effect of central methylation (Figure S6A). However, it failed to quantitatively capture the relative magnitude of these opposing positional effects and could not reproduce the experimentally observed enhanced binding affinities when methylation occurred at both the center and flanking regions. We further tested other training combinations in which the dataset contained sequences with a CpG at the central site, as well as combinations that included both central-CpG and flanking-CpG sequences, yet none yielded satisfactory results (Figures S6B and C). These results suggest that although training with static structures and their associated sequences can capture position-dependent DNA methylation effects, the data remain insufficient to resolve the subtle positional specificities, highlighting the necessity of integrating additional experimental data to improve model sensitivity to methylation context.

**Fig. 3.**
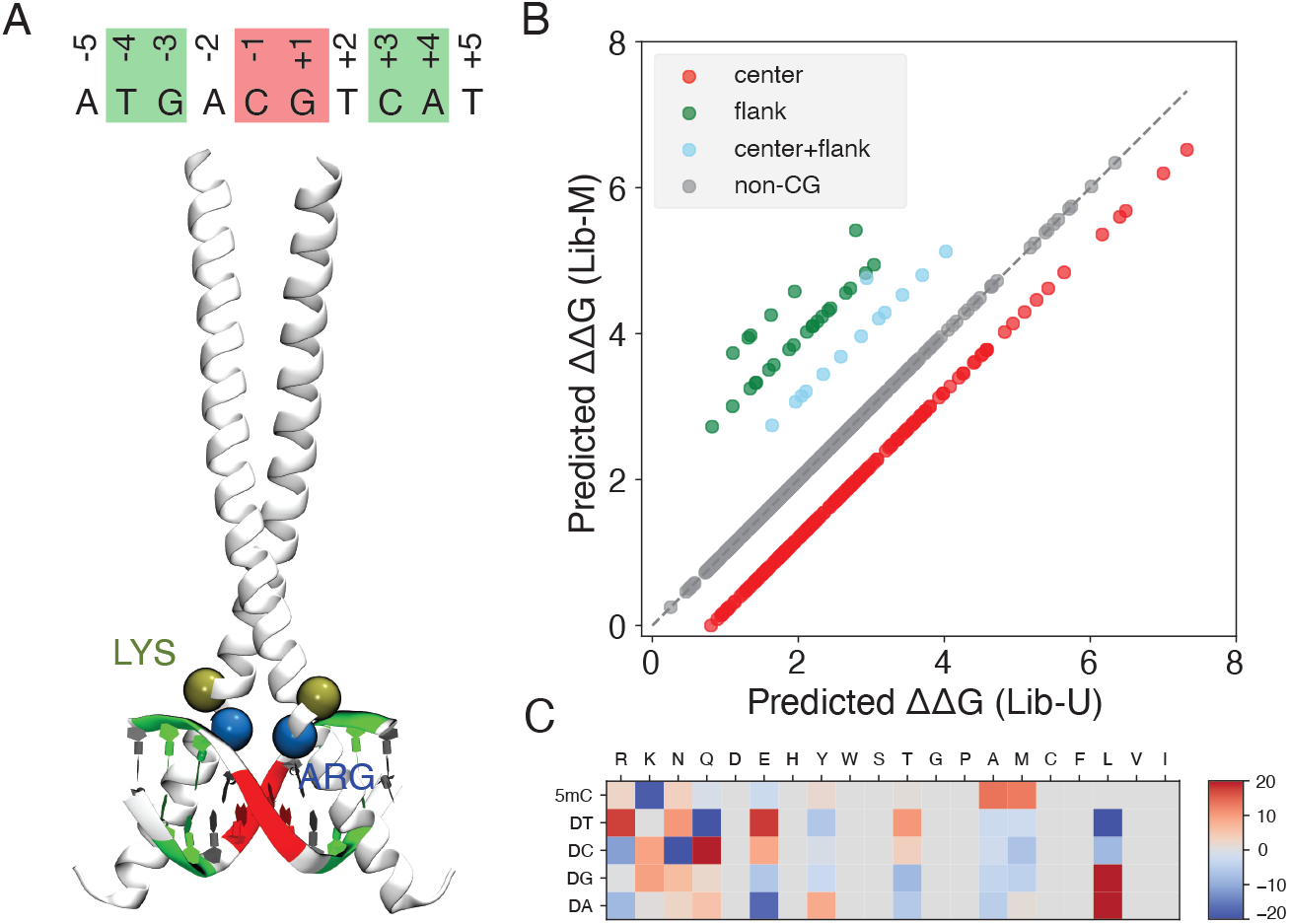
mIDEA decodes position-specific methylation sensitivity of ATF4. (**A**) AF3-predicted structure of the ATF4–DNA complex showing the cyclic AMP response element (CRE) motif (5’-TGACGTCA-3’). (**B**) Predicted binding free energies of ATF4 on methylated (Lib-M) versus unmethylated (Lib-U) 10-mer ligands. Methylation at flanking sites enhances binding affinity (green), whereas central methylation reduces it (red). When both regions are methylated (blue), the overall affinity remains elevated but weaker than flanking methylation alone. (**C**) Trained residue–level interaction energy matrix showing distinct amino acid preferences.

We therefore adopted a data-enhanced training strategy (Section 2.1.3) integrating both structural information and EpiSELEX-seq data, with the corresponding training and testing performance at different regularization strengths illustrated in Figure S7. These two data sources are complementary: structural data provides mechanistic contexts, whereas EpiSELEX-Seq data encodes position-specific methylation sensitivity. This integrative approach substantially improved model performance, with predictions that closely matched experimental observations (Figure 3B). Furthermore, the model correctly distinguished between methylation at different flanking regions, predicting slightly stronger effects at the right flank compared to the left. By comparison, this subtle positional methylation effect was not captured by DeepRec [29], a geometry-aware, sequence-based neural network with more than 1,000 parameters, possibly because the EpiSELEX-seq dataset contains only around 500 high-confidence ligand affinity measurements, which may be insufficient to robustly train a model of this complexity. Therefore, mIDEA accurately recapitulates experimentally observed methylation-dependent effects, suggesting that the AF3-predicted ATF4–DNA complex correctly captures the binding interface of the actual ATF4–methylated DNA structure and demonstrates the benefit of incorporating structural information in predicting protein–DNA binding specificity, particularly when the available training data are not extensive.

The trained energy model (Figure 3C) reveals distinct patterns of interactions between unmodified and methylated cytosines and specific amino acids. Notably, 5mC exhibits favorable interactions with LYS, whereas unmodified cytosine interacts more strongly with ARG. We hypothesize that these dominant residue-specific interactions play a key role in shaping ATF4’s position-dependent response to CpG methylation. Structural analysis of the AF3-predicted complex (Figure 3A) shows that two symmetric LYS residues are positioned in close proximity to the flanking regions, whereas ARG residues are located near the central site. These distinct amino acid–nucleotide interactions thereby provide a mechanistic basis for ATF4’s position-specific sensitivity to CpG methylation.

### 3.4 Methylation-Informed Identification of Genomic Protein-DNA Binding Sites

DNA methylation also plays an important role in shaping the genome-wide binding landscape of regulatory proteins [12,13,14]. High-throughput methods such as ChIP-seq [47,48,49], ChIP-exo [50,51,52], and CUT&RUN [53] have enabled precise mapping of protein–DNA interactions across the genome. Meanwhile, genome-wide DNA methylation profiling can be achieved using whole-genome bisulfite sequencing (WGBS) [54,55], reduced representation bisulfite sequencing (RRBS) [56], and MeDIP-seq[57], which together offer complementary trade-offs between resolution and cost. Build on mIDEA’s performance in predicting *in vitro* binding, we next evaluated its applicability to predict methylation effect in an *in vivo* condition. We focused on a 1 Mb region (156000–157000 kb) on chromosome 1 of GM12878 cell line, which shows the strongest and most reliable MAX binding signals and were previously captured by our sequence-specific IDEA model [36]. Given that MAX is a methyl-minus protein (Figure S3), we expected its binding sites to be depleted of methylated CpGs. Analysis of ChIP-seq and WGBS data confirmed this pattern (Figure 4A), indicating that MAX’s methylation sensitivity is consistent between *in vitro* and *in vivo* contexts. To incorporate methylation information for enhancing predictions of genomic protein binding sites, we implemented a methylation-informed prediction framework (Section 2.2) that weights CpG contributions based on their methylation levels. This approach improved the model’s overall performance, achieving an AUC score of 0.84 and a balanced PRAUC score [46] of 0.83 by reducing false positive predictions while maintaining recovery of true ChIP-seq peaks. This represents an improvement over both the sequence-specific unmethylated model and the model assuming universal CpG methylation (Figure 4B and C).

**Fig. 4.**
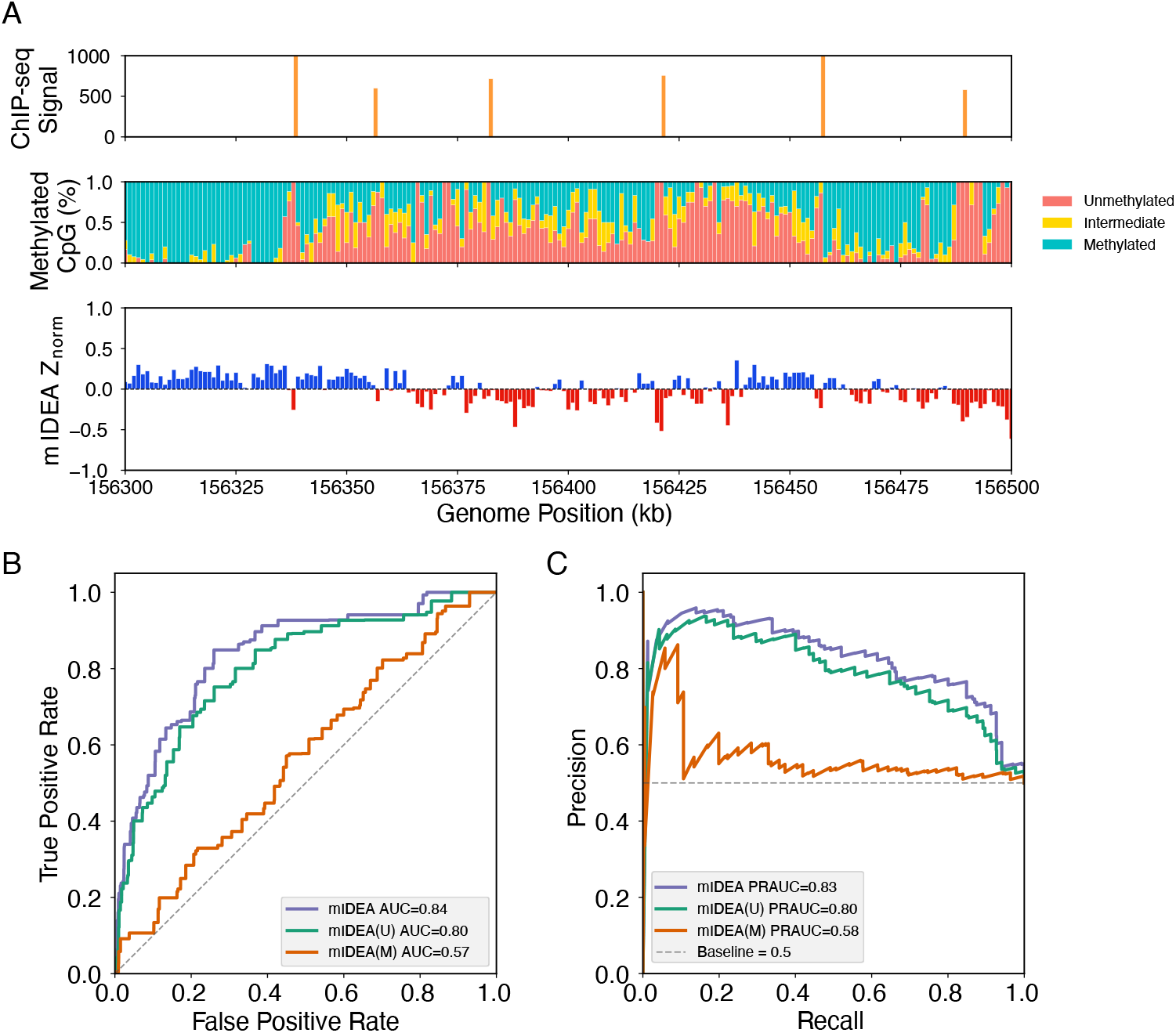
Performance of methylation-informed mIDEA for MAX binding prediction. (**A**) Multi-track visualization of MAX binding landscape in a 200 kb genomic region (chr1:156300–156500 kb). From top to bottom: ChIP-seq signals; (ii) CpG methylation state classification based on WGBS data (red: unmethylated, 30%; yellow: intermediate, ≤30–70%; blue: methylated, ≥70%), (iii) mIDEA-predicted binding energy Z-scores averaged over 1 kb bins (red bars, *Z <* 0, predicted high-confidence binding; blue bars, *Z ≥* 0, predicted unfavorable binding). (**B**) Receiver operating characteristic (ROC) curves comparing mIDEA, mIDEA(U) (sequence-specific version without methylation), and mIDEA(M) (assuming all CpGs are methylated) models, evaluated at 500 bp resolution to match ChIP-seq signal precision, with the Area Under the Curve (AUC) scores are labeled. (**C**) Precision–Recall curves for the same models (500 bp resolution). Balanced precision-recall AUC (PRAUC) scores [46] are labeled, with the dashed line representing random classifier performance (PRAUC = 0.5).

Interestingly, incorporating methylation information does not improve the prediction of methyl-plus proteins such as HOXB13. In A549 cells, HOXB13 binding sites within the 1 Mb region on chromosome 17 (59,000–60,000 kb) exhibit lower methylation levels than their flanking sequences (Figure S8). This pattern suggests that methylation is unlikely to be the primary determinant of HOXB13 binding. Instead, HOXB13 may initially recognize methylated CpG sites and subsequently cause local DNA demethylation[6,15,9]. Nevertheless, the precise mechanism underlying this methylation–binding interplay remains to be elucidated[22].

## 4 Discussion

mIDEA is a structure-based, computationally efficient biophysical framework for investigating the effects of cytosine methylation on protein–DNA affinities. Compared with existing approaches[25,21,26,27,28,29], its key innovation lies in the ability to jointly learn from structural and sequencing data via a unified, residue-level energy model. This structure–sequence integration enables accurate prediction of methylation effects even with limited training data and provides mechanistic insights into how DNA methylation modulates protein–DNA interactions through the learned energy parameters.

Although mIDEA has been successful in modeling DNA methylation, the protocol can be improved in several aspects. The current implementation assumes that both methylated and unmethylated protein–DNA complexes share the same structural interface. However, recent studies using methyl-DNAshape [16] have shown that CpG methylation can significantly alter local DNA structure. To address this, an important next step is to augment the training datasets with high-confidence predicted protein–methylated DNA structures [31,32,33,34] to better capture methylation-induced structural variation.

Moreover, the existing framework for predicting genomic DNA-binding loci considers only DNA sequences and CpG methylation profiles. In addition to sequence features, chromatin accessibility is known to substantially influence protein–DNA binding specificity. Incorporating accessibility information, such as ATAC-seq [58] and DNase-seq [59], into the predictive framework could further improve model accuracy.

Since mIDEA provides sequence- and epigenetic-specific interaction strengths for each amino acid–nucleotide pair, it can be straightforwardly incorporated into molecular dynamics (MD) force fields [36]. Simulations using these integrated force fields could yield deeper understanding of the dynamic and mechanistic basis of methylation-modulated genomic regulation.

Finally, the mIDEA framework can be extended to model protein interactions with other unmodified and modified nucleic acids [36,43], including DNA modifications such as 5-hydroxymethylcytosine (5hmC), 5-formylcytosine (5fC), and 5-carboxylcytosine (5caC), as well as RNA modifications [2] such as N6-methyladenosine (m6A), N6,2’-O-dimethyladenosine (m6Am), N1-methyladenosine (m1A), and 5-methylcytosine (m5C). These chemical modifications can significantly alter nucleic acid stability, protein interactions, and downstream gene regulation [60,61,62].

## Acknowledgments

This work was supported by startup funding from North Carolina State University, with additional support from the NC State Genetics and Genomics Academy and the Comparative Medicine Institute.

