## Supplementary Materials for "mIDEA: An Interpretable Structure–Sequence Model for Methylation-Dependent Protein–DNA Binding Sensitivity"

### S1 Supplementary Texts

#### S1.1 Data Collection

**S1.1.1 Structural Data** The experimental structural data were obtained from the Protein Data Bank (PDB, https://www.rcsb.org/). The PDB IDs for C/EBP$\beta$, HOXB13, and MAX are 8K8D, 5EF6, and 1HLO, respectively.

We also used AlphaFold3 to model a high-confidence ATF4 homodimer–DNA complex. For the input protein sequences, two identical ATF4 monomers were used, derived from the previously crystallized ATF4–C/EBP$\beta$ heterodimer (PDB ID: 1CI6). The DNA sequence used in the prediction was 5'-ACGATGTCAT-3', which corresponds to the top 10-mer binding motif identified from the EpiSELEX data.

**S1.1.2 Sequencing Data** The EpiSELEX-seq data, containing the relative affinities and corresponding oligomer sequences for C/EBP$\beta$ and ATF4 in methylated (Lib-M) and unmethylated (Lib-U) libraries, were obtained from the NCBI Gene Expression Omnibus (GEO: GSE98652).

The methyl-HT-SELEX data for HOXB13 and MAX were retrieved from the European Nucleotide Archive (ENA: PRJEB9797). We primarily used the R package `SELEX` (https://www.bioconductor.org/packages/release/bioc/html/SELEX.html) to process rounds 1 and 4 of both the normal SELEX and methyl-SELEX experiments to derive k-mer counts and relative binding affinities. To ensure data quality, we filtered the ligands by retaining only those with more than 200 counts in both libraries.

The ChIP-seq data for HOXB13 in the GM12878 cell line were obtained from the ENCODE project (accession ID: ENCFF361EVH), and the corresponding whole-genome bisulfite sequencing (WGBS) data were retrieved under accession ID: ENCFF279HCL. For the A549 cell line, the ChIP-seq data for HOXB13 were obtained from ENCODE (accession ID: ENCSR967ZMR), and the corresponding WGBS dataset was accessed under accession ID: ENCFF003JVR. The reference human genome (GRCh38) was downloaded from the UCSC Genome Browser at https://hgdownload.soe.ucsc.edu/goldenPath/hg38/chromosomes/.

### S2 Supplementary Figures

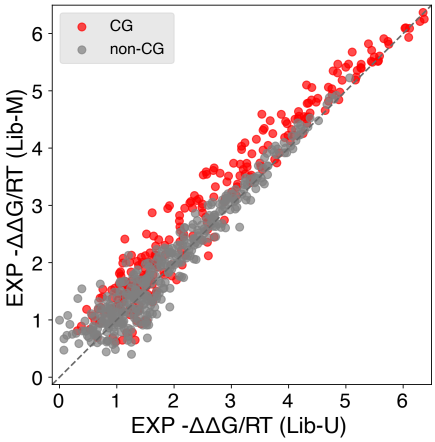


**Fig. S1. Experimental observation of the weak methyl-plus binding effect of C/EBP$\beta$.** Experimental binding free energies (EXP $-\Delta\Delta G/RT$ ) of C/EBP$\beta$ derived from EpiSELEX-seq, comparing methylated (Lib-M) and unmethylated (Lib-U) libraries. Each point represents a 10-mer ligand; CG-containing sequences (red) exhibit slightly higher binding affinities in the methylated library, consistent with a weak methyl-plus effect, whereas non-CG sequences (gray) remain largely unchanged.

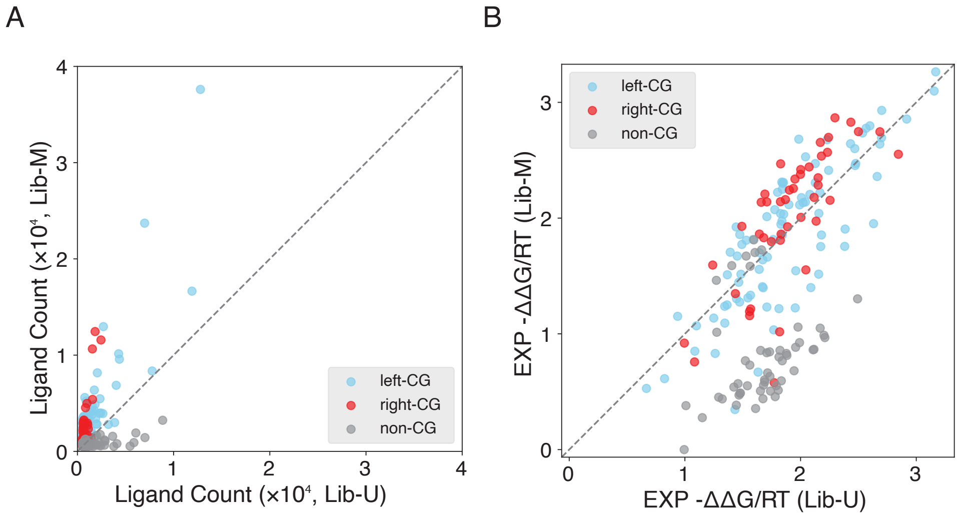


**Fig. S2. Experimental evidence showing that HOXB13 exhibits strong methyl-plus binding effects at right-flanking mCpG sites.** (**A**) Ligand counts of 10-mer sequences from methylated (Lib-M) and unmethylated (Lib-U) libraries, showing enrichment of CG-containing sequences, particularly those with right-flanking CpG sites. (**B**) Experimental binding free energies derived from methyl-HT-SELEX, revealing that methylation at right-CG sites generally enhances binding affinity, whereas left-CG methylation exhibits variable effects.

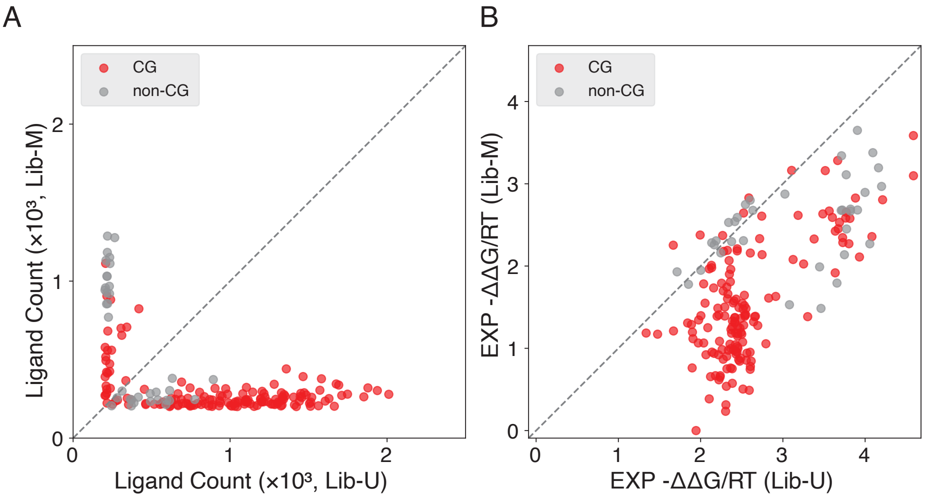


**Fig. S3. Experimental evidence showing that MAX exhibits methyl-minus binding behavior at CpG sites.** (**A**) Ligand counts of 11-mer sequences from methylated (Lib-M) and unmethylated (Lib-U) libraries, showing that CpG-containing ligands are markedly depleted upon methylation, consistent with reduced enrichment in the methylated library. (**B**) Experimental binding free energies derived from methyl-HT-SELEX, further demonstrating that CpG methylation generally weakens binding affinity for MAX compared to the unmethylated counterparts, reflecting its strong methyl-minus specificity.

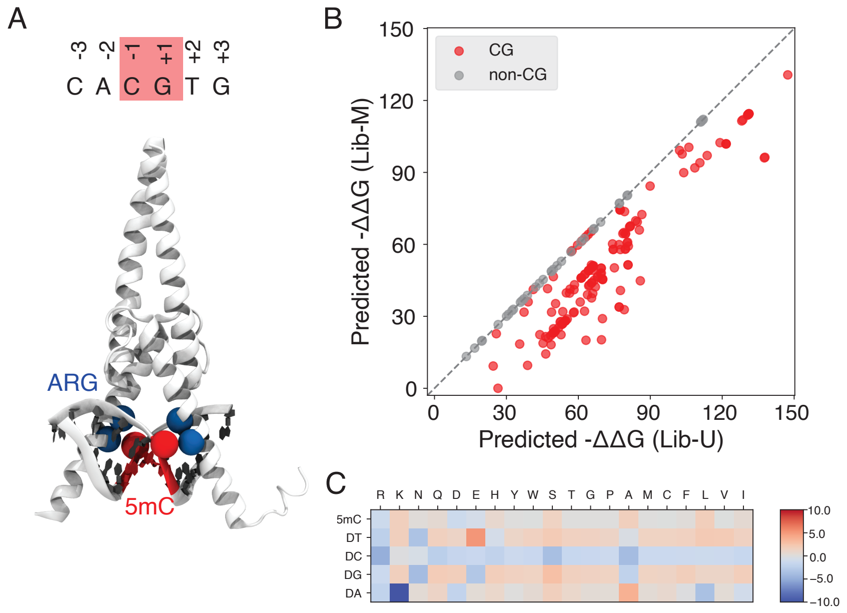


**Fig. S4. Modeling methyl-minus binding specificity of MAX using mIDEA.** (**A**) Crystal structure of the MAX–DNA complex (PDB ID: 1HLO), highlighting the E-box core motif (5'-CACGTG-3') and the central CpG dinucleotide that serves as the primary methylation site. (**B**) Predicted binding free energies for methylated (Lib-M) versus unmethylated (Lib-U) ligands. Most ligands are methylated at the central CG of the E-box motif and show reduced binding affinity upon CpG methylation, consistent with the experimentally observed methyl-minus behavior. (**C**) Trained residue–base energy map, showing stronger ARG–cytosine interactions compared to ARG–5mC, revealing the molecular basis for MAX's reduced affinity toward methylated DNA.

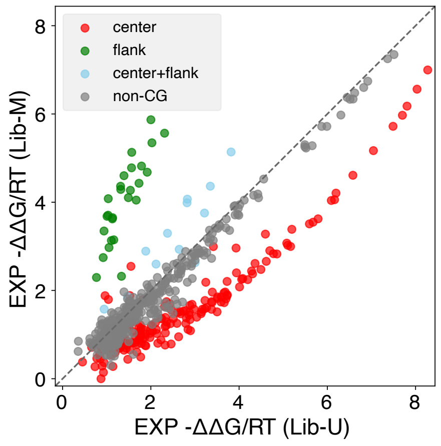


**Fig. S5. Experimental evidence showing position-specific methylation effects on ATF4 binding.** EpiSELEX-seq–derived binding free energies for methylated (Lib-M) versus unmethylated (Lib-U) ligands. Methylation at flanking CpG sites (green) markedly enhances ATF4 binding, whereas methylation at the central CpG (red) reduces it. Double methylation at both center and flanks (blue) produces an intermediate effect, confirming that flanking methylation dominates the net affinity change.

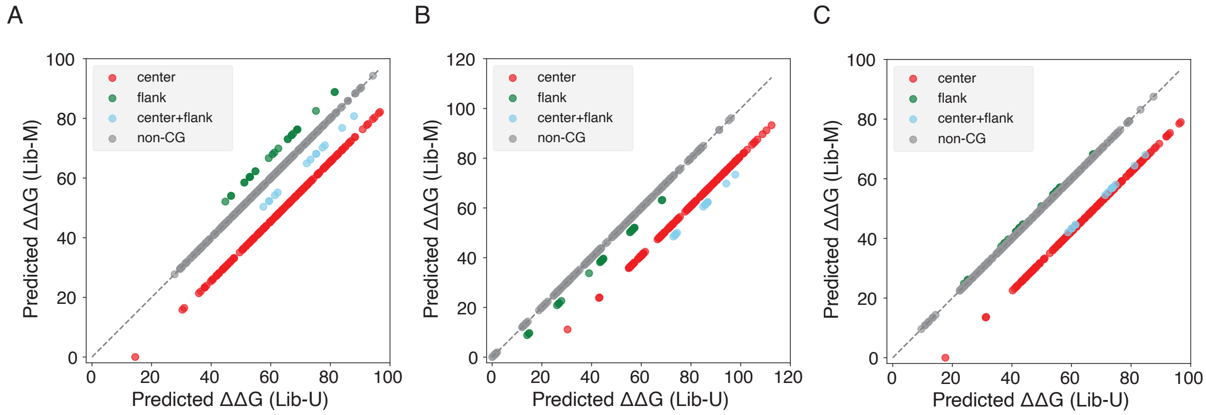


**Fig. S6. Testing three training combinations for modeling ATF4 methylation effects using strategy I.** (**A**) The mIDEA model was trained using a 10:1 ratio of methylated to unmethylated left-flank CG sequences (5'-ACGATGTCAT-3') to emphasize the dominant flank-methylation effect. (**B**) The model was trained using sequences containing CpG at the central position (5'-ATGACGTCAT-3') with an inverse 1:10 methylated-to-unmethylated ratio to capture the methylation-suppressive effect at the center. (**C**) The model was trained using a training set composed of a mixture of both flanking and central methylation templates.

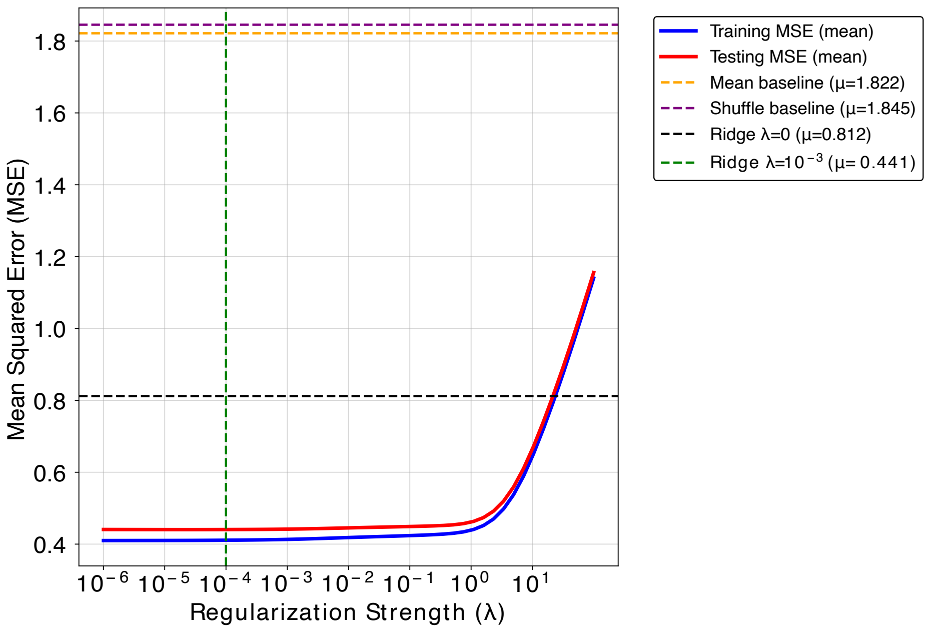


**Fig. S7. Optimization of regularization strength for fitting ATF4 binding energies using ridge regression.** Experimental binding energies ($E_{\text{exp}}$) of ATF4–DNA complexes were fitted using ridge regression across a range of regularization strengths $\lambda$ from $10^{-6}$ to $10^{2}$. For each $\lambda$ value, performance was evaluated by averaging results over 50 independent train–test splits (80% training, 20% testing). Blue and red curves represent the mean training and testing mean squared errors (MSE), respectively, as functions of $\lambda$ on a logarithmic scale. The black dashed line denotes the performance of the unregularized least-squares model ($\lambda = 0$). The dashed green vertical line marks the selected regularization strength, $\lambda = 10^{-4}$, located within the flat minimum region of the test-error curve. This value was used to refit the final ridge model on the complete dataset. To evaluate the contribution of sequence-specific training energies, we compared mIDEA's performance against baseline models that lack sequence specificity. Specifically, we constructed (i) a mean-energy baseline that assigns a uniform average energy to all sequences (orange dashed line), and (ii) a shuffle baseline in which experimental energies were randomly permuted across sequences before training (purple dashed line). For both baselines, energy matrices were derived using the corresponding non–sequence-specific energies, and MSEs were averaged over all random splits. The substantially higher baseline MSEs in both cases highlight the importance of the sequence-dependent interaction strengths learned by mIDEA.

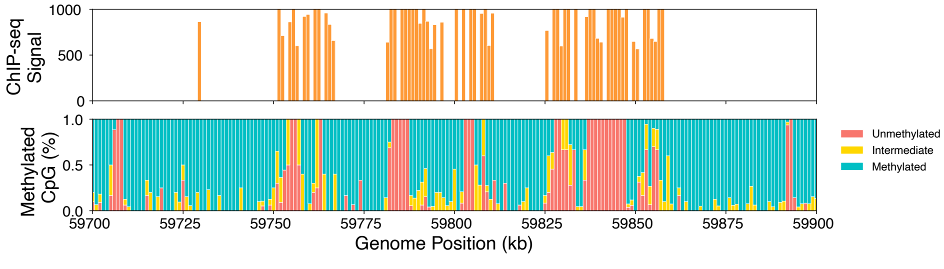


**Fig. S8. Methylation-aware genomic analysis of HOXB13 binding landscape.** Two-track genomic view showing ChIP-seq signal (top) and CpG methylation levels (bottom) across a 200 kb region on chromosome 17 (59700–59900 kb) in A549 cells. Although HOXB13 is characterized as a methyl-plus protein, its binding sites in this region exhibit locally reduced methylation relative to the surrounding regions, suggesting either a divergence between *in vitro* and *in vivo* binding preferences or a methylation-dependent recruitment process followed by localized demethylation.
